# Temperature-responsive competitive inhibition of CRISPR-Cas9

**DOI:** 10.1101/407718

**Authors:** Fuguo Jiang, Jun-Jie Liu, Beatriz A. Osuna, Michael Xu, Joel D. Berry, Benjamin J. Rauch, Eva Nogales, Joseph Bondy-Denomy, Jennifer A. Doudna

## Abstract

CRISPR–Cas immune systems utilize RNA-guided nucleases to protect bacteria from bacteriophage infection. Bacteriophages have in turn evolved inhibitory ‘anti-CRISPR’ (Acr) proteins, including six inhibitors (AcrIIA1-6) that can block DNA cutting and genome editing by type II-A CRISPR-Cas9 enzymes. We show here that AcrIIA2 and its homologue, AcrIIA2b, prevent Cas9 binding to DNA by occluding protein residues required for DNA binding. Cryo-EM-determined structures of AcrIIA2 or AcrIIA2b bound to *S. pyogenes* Cas9 reveal a mode of competitive inhibition of DNA binding that is distinct from other known Acrs. Differences in the temperature dependence of Cas9 inhibition by AcrIIA2 and AcrIIA2b arise from differences in both inhibitor structure and the local inhibitor-binding environment on Cas9. These findings expand the natural toolbox for regulating CRISPR-Cas9 genome editing temporally, spatially and conditionally.

## INTRODUCTION

Bacteriophage are the most abundant biological entity on the planet and impart strong selective pressure on their bacterial hosts. In addition to innate defense systems, bacteria have developed adaptive immunity known as CRISPR–Cas to recognize and destroy foreign nucleic acids in a sequence-specific manner (Barrangou and Marraffini, 2014; Hille and Charpentier, 2016; Marraffini, 2015; 2010). CRISPR–Cas systems are classified into six diverse types (I-VI) (Koonin et al., 2017; Makarova et al., 2015) that use a CRISPR genomic sequence array to record genetic evidence of prior infections. Small RNA guides transcribed from the array, together with Cas nucleases, target and degrade phage DNA or RNA (Hale et al., 2009; Marraffini and Sontheimer, 2008; Wiedenheft et al., 2011).

To counteract CRISPR-Cas immunity, phage employ inhibitory proteins to inactivate CRISPR–Cas function in a sequence-independent manner (Bondy-Denomy et al., 2013; Sontheimer and Davidson, 2017). To date, >20 diverse anti-CRISPRs have been identified in phage, prophage and mobile genetic elements (Borges et al., 2017). Notably, four distinct anti-CRISPR proteins that inhibit type II-A CRISPR-Cas9 (AcrIIA1-4) from *Listeria monocytogenes* prophage were identified along with three that inactivate type II-C Cas9 orthologs (AcrIIC1-3), representing the first identification of anti-CRISPR proteins in type II CRISPR–Cas systems (Pawluk et al., 2016; Rauch et al., 2017). More recently, AcrIIA5 and AcrIIA6 has also been discovered in *Streptococcus thermophilus* phages (Hynes et al., 2018; 2017). Two of these inhibitors, AcrIIA2 and AcrIIA4, possess a broad-spectrum host range by inhibiting the activity of *Streptococcus pyogenes* Cas9 (53% amino acid identity to *L. monocytogenes* Cas9) in bacterial and human cells, although the ability of AcrIIA2 to block Cas9 functions is weaker than that of AcrIIA4 (Rauch et al., 2017). AcrIIA4 can function as a gene editing “off-switch” in human cells by reducing off-target mutations (Shin et al., 2017), by limiting Cas9-mediated toxicity in hematopoietic stem cells (Li et al., 2018) and by halting dCas9-based epigenetic modifications (Liu et al., 2018). Additionally, AcrIIA2 and AcrIIA4 have been used to limit Cas9-mediated gene drives in yeast (Basgall et al., 2018), demonstrating wide-ranging utility for these proteins. Structural studies showed that AcrIIA4 acts as a DNA mimic and binds to the PAM-interacting motif of the Cas9 protein to prevent target DNA binding and cleavage (Dong et al., 2017a; Shin et al., 2017; Yang and Patel, 2017). Biochemical work suggested that AcrIIA2 might act in a similar manner (Dong et al., 2017a; and Yang and Patel, 2017). However, the structural basis of AcrIIA2-mediated inhibition of Cas9–DNA binding remains obscure.

To determine the mechanism of AcrIIA2-mediated Cas9 inhibition, and to explore its utility as an effective “off-switch” for CRISPR–Cas9 regulation in genome editing applications, we determined a 3.4 Å resolution cryo-EM structure of AcrIIA2 interacting with sgRNA-loaded SpyCas9. Additionally, we identified a homolog of AcrIIA2 (AcrIIA2b.3), encoded on an *L. monocytogenes* plasmid, which has more robust SpyCas9 inhibitory activity both *in vitro* and *in vivo*. A 3.9-Å cryoEM structure of AcrIIA2b.3 bound to SpyCas9 revealed a binding pocket similar to that observed in AcrIIA4 for blocking PAM recognition which results in a more robust inhibition by AcrIIA2b.3 relative to AcrIIA2. We show that temperature dependent inhibition occurs *in vitro* and likely results from differences in inhibitor stability at different temperatures. This work provides a comprehensive analysis of CRISPR–Cas9 functional interference mediated by the AcrIIA2 inhibitor family, but also a framework for future structure-based anti-CRISPR engineering and small peptide inhibitor design for precise and efficient control of Cas9-mediated genome editing.

## RESULTS

### Architecture of AcrIIA2 bound to sgRNA-loaded SpyCas9

AcrIIA2 is a type II-A anti-CRISPR commonly found in phages and prophages of *L. monocytogenes*, comprising 123 amino acids, that inhibits SpyCas9 both *in vitro* and *in vivo* (Basgall et al., 2018; Rauch et al., 2017; Yang and Patel, 2017). We first investigated at which step of CRISPR–Cas9 assembly AcrIIA2 inactivates Cas9 function. We performed size-exclusion chromatography (SEC) to test whether AcrIIA2 physically interacts with either SpyCas9 or sgRNA, or with the binary complex. Consistent with previous biochemical observation(Yang and Patel, 2017), AcrIIA2 can only form a stable complex with sgRNA-loaded SpyCas9, and no direct interaction occurs with either apo-SpyCas9 or sgRNA alone (Figures 1 and S1A–1C). This chromatographic profile of complex formation is similar to what has been observed for AcrIIA4 (Dong et al., 2017b; Shin et al., 2017; Yang and Patel, 2017), indicating that AcrIIA2 interacts with SpyCas9 in a similar manner to AcrIIA4 and most likely binds to Cas9 at the region that is created upon sgRNA-triggered conformational rearrangement in Cas9 (Jiang et al., 2015).

**Figure 1.**
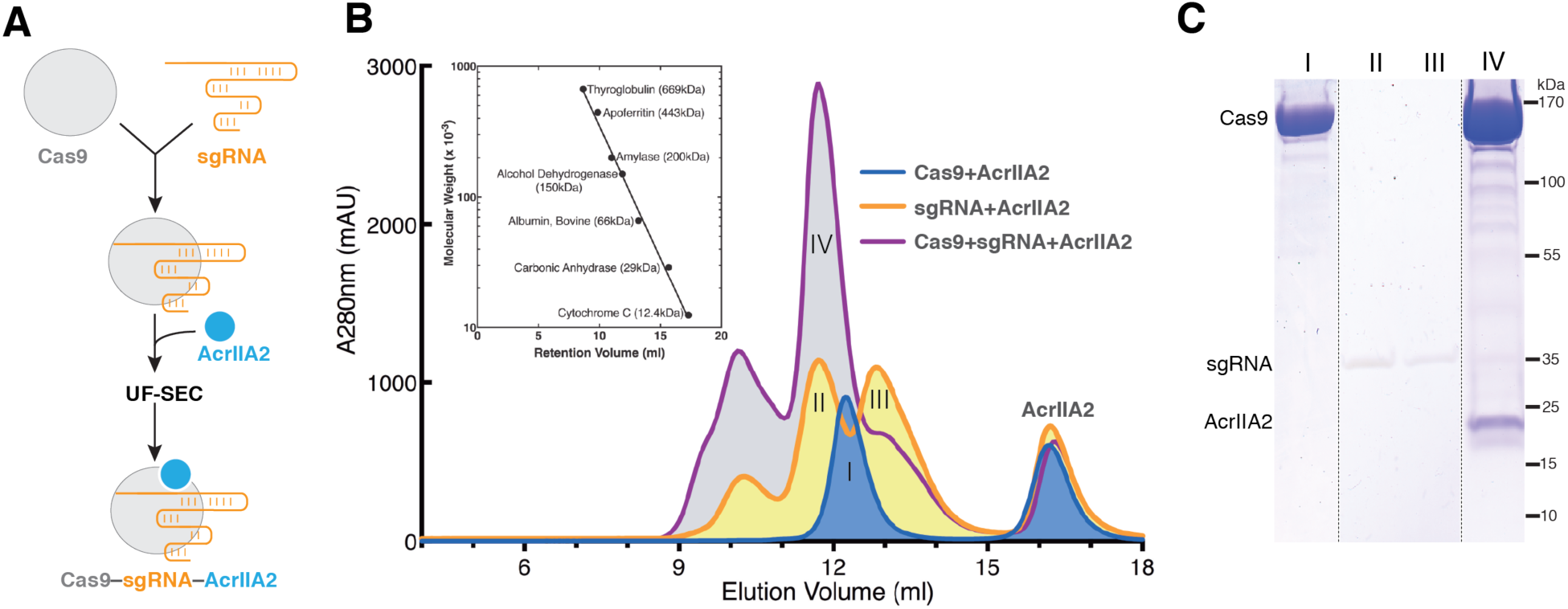
AcrIIA2 selectively forms a stable complex with the sgRNA-loaded *S. pyogenes* Cas9 rather than apo- or DNA-bound SpyCas9. (A). Flow chart for reconstitution and isolation of the AcrIIA2-bound SpyCas9–sgRNA ternary complex using ultrafiltration combined with size exclusion chromatography (UF-SEC). (B) Size exclusion chromatogram of SpyCas9–AcrIIA2 in the presence or absence of sgRNA. The inset represents the calibration curve obtained using standard molecular weight markers. (C) Coomassie stained polyacrylamide gel showing the co-purification of AcrIIA2 with sgRNA-bound SpyCas9.

In order to elucidate the detailed structural mechanism of AcrIIA2-mediated inhibition of Cas9 activity, we obtained images of the frozen-hydrated samples (Figure S2A), and solved the cryo-EM structure of SpyCas9–sgRNA–AcrIIA2 complex with an overall resolution of 3.4 Å (Figures 2, S2–S5, and Table S1). We observed electron density corresponding to the AcrIIA2 inhibitor protein with clear side chain features (Figure 2A–B) that enabled atomic modeling of AcrIIA2 (Figure 2C). In contrast to the well-defined complex core region and the bound AcrIIA2, the HNH domain within the SpyCas9 shows weaker density (Figures S5 and S6D), reflecting the intrinsic flexibility of this nuclease domain, as seen in prior structures (Anders et al., 2014; Jiang et al., 2015; 2016; Nishimasu et al., 2014).

**Figure 2.**
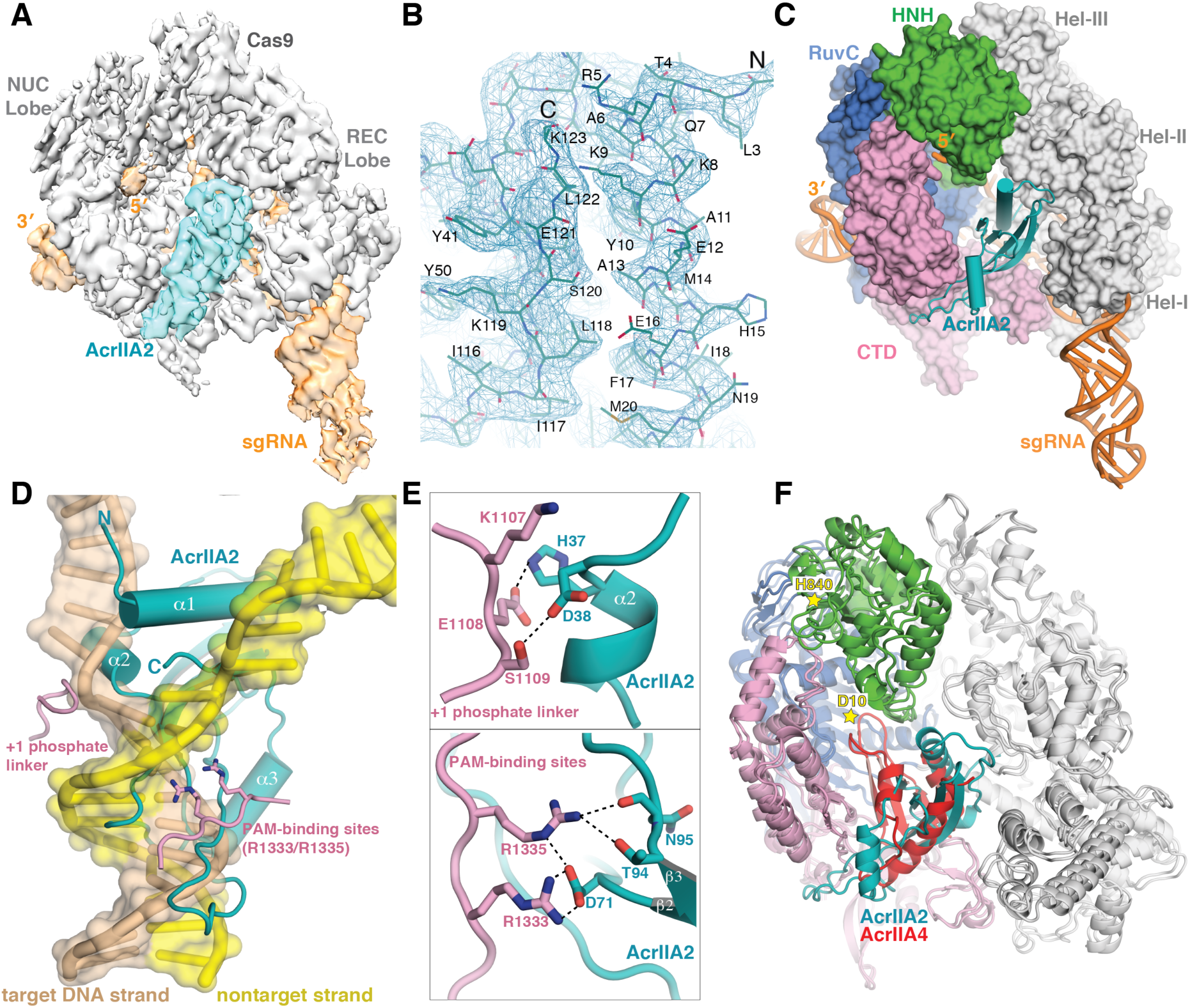
Architecture of the *S. pyogenes* Cas9–sgRNA in complex with AcrIIA2. (A) Cryo-EM reconstruction of SpyCas9–sgRNA–AcrIIA2 at 3.4-Å resolution, segmented to highlight densities corresponding to SpyCas9 (gray) and AcrIIA2 (teal). (B) Representative cryo-EM density for AcrIIA2 with the refined model superimposed. (C) Surface representation of SpyCas9–sgRNA–AcrIIA2 ternary complex. (D) Superimposition of AcrIIA2 (teal) with the dsDNA-bound SpyCas9 complex. For clarity, Cas9 is omitted except the target DNA strand and non-target strand. (E) Close-up view showing how AcrIIA2 blocks target DNA binding by interacting with SpyCas9’s PAM recognition residues (R1333/R1335) and +1 phosphate linker. (F) Superposition with SpyCas9–sgRNA–AcrIIA4 structure (PDB ID 5VZL). Stars indicate the two catalytic residues (D10 of RuvC and H840 of HNH).

In the refined atomic model of the SpyCas9–sgRNA–AcrIIA2 complex, the conformation of SpyCas9 resembles the pre-target-bound state rather than dsDNA-bound state (Figure S6A– C). This suggests that AcrIIA2 binding blocks Cas9 conformational rearrangement, especially the HNH catalytic domain movement from the inactive state to the active state. Our *ab initio* modeling of the AcrIIA2-bound SpyCas9 complex demonstrated that AcrIIA2 sits within the nucleic-acid-binding channel that forms only upon sgRNA loading between the nuclease lobe (NUC) and helical recognition lobe (REC). This observation explains why sgRNA loading is critical for AcrIIA2 binding to Cas9. AcrIIA2 binds to the SpyCas9–sgRNA complex with 1:1 stoichiometry and at the same binding location as AcrIIA4 (Figures 2C and 2F). It is also worth noting that AcrIIA2 exists as a monomer in solution and maintains a single domain structure in the complex.

The structure of AcrIIA2 is a mixed α+β fold, composed of a bent four-stranded antiparallel β sheet with a β4β1β3β2 arrangement, flanked by two helices, one on each side (Figure S7B–C). Notably, this topology of AcrIIA2 (α1α2β1β2β3α3β4) is distinct from that of AcrIIA4 (α1β1β2β3α2α3) and any other reported anti-CRISPR structures (Figure S7H–I). AcrIIA2 is structurally similar to aspartate-kinase chorismate-mutase tyrA (ACT) (Figure S8), a regulatory domain found in a variety of proteins that exhibits low sequence conservation and high functional divergence from AcrIIA2 (Grant, 2006).

### AcrIIA2 directly blocks target DNA binding

Based on the cryo-EM structure of the AcrIIA2–bound SpyCas9 complex, we analyzed the detailed interactions between Cas9 and AcrIIA2. Superposition of the AcrIIA2-bound SpyCas9 structure onto the dsDNA-bound SpyCas9 structure reveals that AcrIIA2 is anchored into the PAM duplex interacting cleft to preclude target DNA binding to Cas9 (Figures 2D and S7A). Close inspection shows that AcrIIA2 likely makes an elaborate intermolecular hydrogen bonding network with the PAM-recognition site (R1333 and R1335) located in Cas9’s C-terminal domain, primarily through the side chains of D71, T94 and the main chain of N95 (Figure 2D–E). Consistent with this observation, the electrostatic potential distribution of AcrIIA2 reveals a highly negative pocket for interaction with the electropositive PAM-recognition site (Figure S9E). In addition, the surrounding residues S1136 and S1338, which are important for recognition of the PAM-proximal DNA duplex in the dsDNA-bound complex, are instead recognized by residues D96 and D60 in the AcrIIA2-bound structure (Figure S7A–B). Given the fact that PAM recognition is the first and key step for target DNA binding and unwinding (Sternberg et al., 2014), it is evident that interfering with the PAM-binding sites deployed by AcrIIA2 (as also seen in AcrIIA4) is an effective means to abolish Cas9-mediated target DNA binding and cleavage activities. To test whether PAM-recognition blockage confers SpyCas9 inhibition by AcrIIA2, we generated single (D71A) or double mutations (T94A/N95A) of AcrIIA2 residues involved in intermolecular contacts with the PAM-recognition site (R1333 and R1335) and analyzed their impact on inhibition of SpyCas9’s *in vivo* DNA targeting activity. These mutations reduced phage survival in the presence of AcrIIA2, with D71A showing essentially complete loss of Cas9 inhibition (Figure 3A). This finding is consistent with *in vivo* evidence showing the loss of gene drive inhibition by unbiased alanine scanning mutagenesis of AcrIIA2 (D65A/D71A) (Basgall et al., 2018).

**Figure 3.**
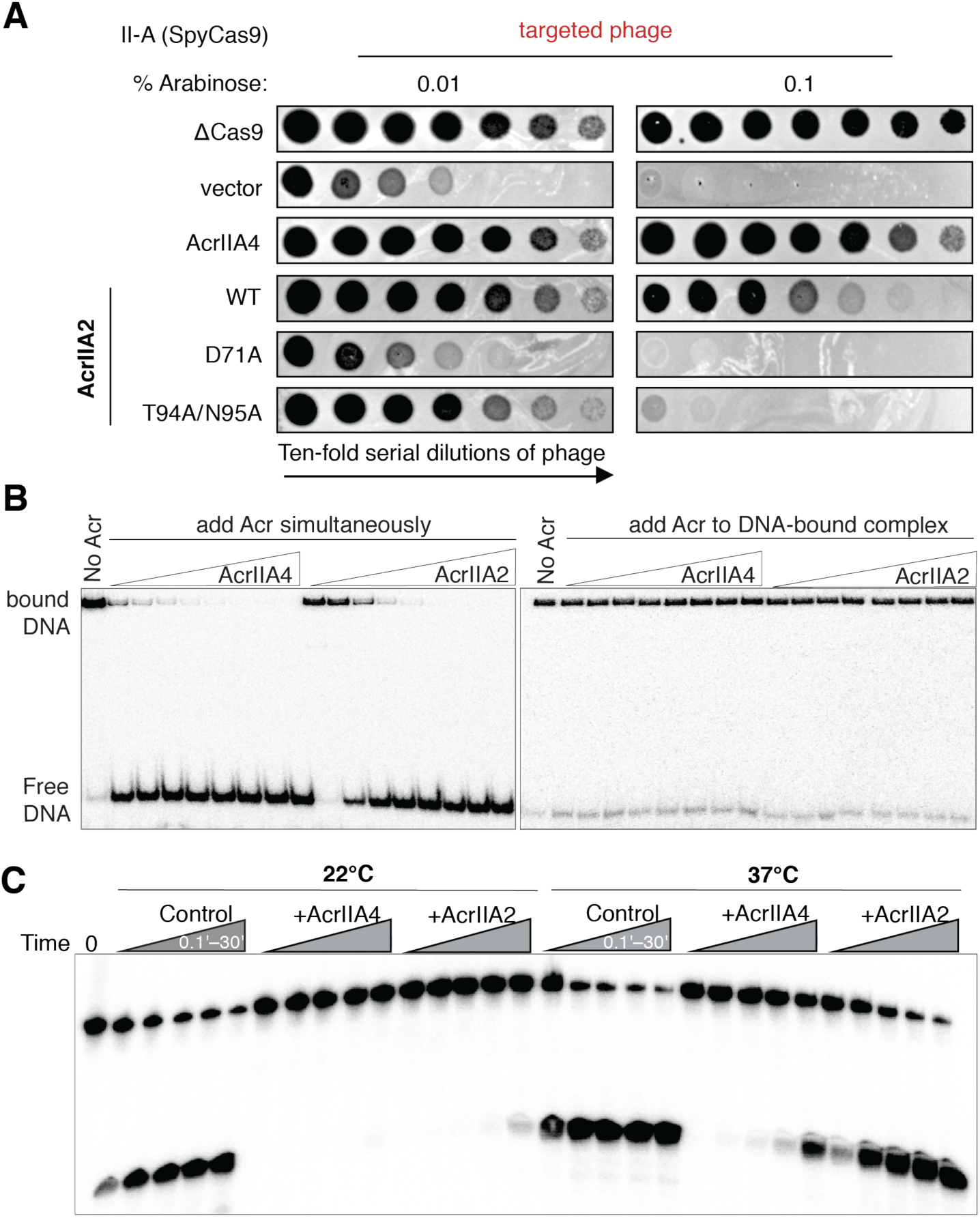
AcrIIA2 is a much less effective inhibitor compared to AcrIIA4. (A) Phage-plaquing experiment where *P. aeruginosa* DMS3m-like phage is titrated in ten-fold dilutions (black circles) on a lawn of *P. aeruginosa* (hazy background) expressing the indicated anti-CRISPR proteins and intermediate (0.01% arabinose) or high (0.1% arabinose) levels of a type II-A CRISPR–Cas9 system programmed to target phage DNA. Plaquing of *P. aeruginosa* phages targeted by SpyCas9 in the presence of AcrIIA2 mutants reveals a nearly complete inactivation of AcrIIA2’s inhibitory effect on SpyCas9 by mutation of D71A. (B) Competition EMSA assays showing AcrIIA2 efficiently competes with target DNA for binding to SpyCas9–sgRNA complex (left), but not the pre-formed DNA-bound complex (right). (C) Radiolabeled cleavage assays conducted using purified SpyCas9–sgRNA complex to assess AcrIIA2 capacity for inhibiting cleavage of both target and non-target DNA strands by SpyCas9 at room temperature and normal body temperature.

In addition to interactions with the PAM-binding site, AcrIIA2 makes extensive contacts with the Helical domain I & II and the other parts of the C-terminal domain from SpyCas9 (Table S2). Specifically, residues H15, N19, E26, and T28 in the N-terminal α1–α2 of AcrIIA2 make hydrophilic interactions with the Helical domain I and domain II. Moreover, the side chains of H37 and D38 of the bound AcrIIA2 interact with the phosphate lock loop (K1107–S1109) located in SpyCas9’s PAM-interacting domain (Figure 2D–E). Previous structural studies indicated that this phosphate lock loop is critical for target DNA unwinding at +1 phosphate on the target DNA strand immediately upstream of the PAM motif (Anders et al., 2014). Superimposing dsDNA-bound and AcrIIA2-bound SpyCas9 reveals that AcrIIA2’s α1 helix wedges between the target strand and nontarget DNA strand, whereas α2 helix penetrates the interface between +1 phosphate lock loop and target DNA strand (Figure 2D). These structural observations indicate that AcrIIA2 binding may also disrupt the target DNA unwinding activity of CRISPR–Cas9 in addition to blocking DNA binding. Indeed, mutational substitution of these residues (D38A/D40A) has demonstrated a large reduction in effectiveness of AcrIIA2 as a Cas9 gene drive inhibitor (Basgall et al., 2018). Apart from these structural features, it is also noteworthy that AcrIIA2 does not interfere with the RuvC active site as observed in AcrIIA4-bound structure (Figure 2F).

We tested and confirmed that AcrIIA2 abrogates SpyCas9-DNA binding by performing gel shift competition assays. Wild-type SpyCas9–sgRNA was used together with 10 mM EDTA to prevent target DNA cleavage. As anticipated, the competition binding experiment with AcrIIA2 and target DNA added simultaneously to SpyCas9-sgRNA resulted in an attenuation or elimination of the dsDNA-bound SpyCas9-sgRNA ternary complex in a concentration dependent manner (Figure 3B). Furthermore, AcrIIA2 showed a weak competitive binding effect compared to AcrIIA4 (Figure S10A–B), indicating that AcrIIA2 binds sgRNA-loaded SpyCas9 with lower affinity than AcrIIA4. Consistent with this, AcrIIA4 can replace AcrIIA2 from the pre-formed anti-CRISPR complex by gel filtration analysis (Figure S11), but not vice versa. Notably, neither AcrIIA2 nor AcrIIA4 bound to dsDNA-bound SpyCas9 (Figures 3B and S11), suggesting that these proteins work *in vivo* by binding Cas9 before the DNA search process is completed (Shin et al., 2017).

### AcrIIA2 is a less effective inhibitor than AcrIIA4

Structural comparison reveals that binding of AcrIIA2 or AcrIIA4 to sgRNA-loaded SpyCas9 results in a similar conformational change within SpyCas9 (Figures 2F and S6D). Moreover, AcrIIA2 sits in the same DNA-binding cavity as AcrIIA4 (Figures S9A and S9G), although they do not bear sequence or structural similarity. In particular, the local environment of AcrIIA2 involved in blocking the PAM-recognition site is almost identical to that of AcrIIA4, except for the lack of two bulky aromatic amino acids (Y41 and Y67 in AcrIIA4) (Figures S7B, S7H, and S9A). Notably, residue Y67 of AcrIIA4 participates in the hydrogen bond network within the PAM-recognition site of SpyCas9, while Y41 forms hydrophobic and van der Waals interactions. We speculate that lack of these bulky hydrophobic residues may result in the lower effectiveness of AcrIIA2-mediated Cas9 inhibition as seen *in vivo*.

We then compared the Cas9-inhibitory activity of AcrIIA2 and AcrIIA4 by *in vitro* DNA cleavage assay. Previous *in vivo* studies showed that AcrIIA4 can completely inactivate Cas9 function, whereas AcrIIA2 only partially blocks Cas9 activity (Rauch et al., 2017). In line with this finding that AcrIIA2 and AcrIIA4 inhibit Cas9 function to different extents, *in vitro* experiments demonstrated that both anti-CRISPR proteins inhibit SpyCas9 enzymatic cleavage activity in a dose-dependent manner (Figures 3C and S12), with AcrIIA2 exhibiting a slightly weaker Cas9 inhibition compared to AcrIIA4 at room temperature (22°C). With large stoichiometric excess of inhibitors over SpyCas9, both AcrIIA2 and AcrIIA4 could fully block Cas9 function (Figure S12), in contrast with previous *in vivo* observations that AcrIIA2 only partially blocks Cas9 function in *coli* and human cells. To resolve this contradiction, we measured AcrIIA2- and AcrIIA4-mediated Cas9 cleavage inhibition at body temperature (37°C). While both inhibitors displayed a substantially decreased level of Cas9 inhibition compared to that observed at lower temperature (Figure 3C), AcrIIA2 failed to inhibit Cas9 even at extreme excess levels (1000:1) (Figure S12G–H). By contrast, a large excess of AcrIIA4 (100:1) can lead to a complete loss of SpyCas9 cleavage activity (Figure S12C–D). Addition of urea to these reactions showed that AcrIIA2-mediated SpyCas9 inhibition is more susceptible to urea-induced denaturation (Figure S13A). Together with results from heat-induced denaturation of Cas9 inhibitors (Figure S14), these data indicate that the AcrIIA2–Cas9 interaction possesses lower thermal stability than the AcrIIA4–Cas9 interaction, which could potentially limit its adoption in control of Cas9-based genome editing.

### Identification of AcrIIA2 homologs with enhanced inhibition activity

Because of the relative inability of the AcrIIA2 protein to inhibit SpyCas9 at human body temperature, we considered whether homologs of *acrIIA2* might possess enhanced inhibition activity against SpyCas9. Homologs of *acrIIA2* are found in prophage, phage, plasmids, and mobile islands within the *Listeria* genus. We identified two distinct sequence families, denoted AcrIIA2b and AcrIIA2c, which possess ∼35% and ∼50% sequence identity, respectively, to the original protein, AcrIIA2 (which corresponds to AcrIIA2a.1) (Figure 4A and S15). To test the function of identified homologs, we established an assay in the bacterium *Pseudomonas aeruginosa* in which a chromosomal copy of SpyCas9 (arabinose-inducible) is programmed to target a *P. aeruginosa* phage, JBD30 (Figure 4B). Candidate *acrIIA2* orthologs were expressed from a plasmid (IPTG-inducible), allowing independent titration of the SpyCas9–sgRNA complex to assess anti-CRISPR strength.

**Figure 4.**
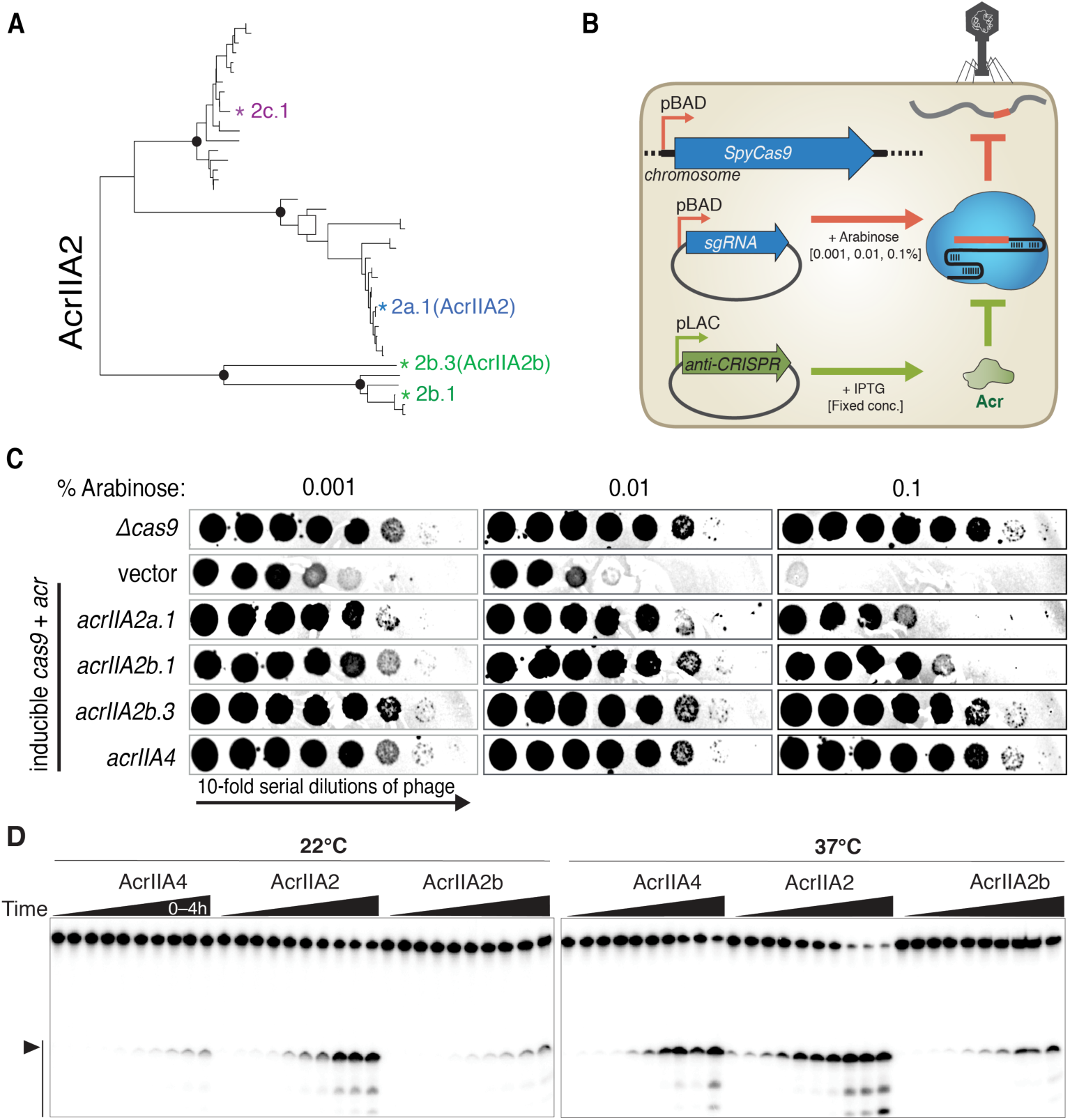
Identifying AcrIIA2b.3 as a potent SpyCas9 inhibitor. (A) Phylogenetic tree of the protein sequences of AcrIIA2 homologs. (B) Schematic of the *P. aeruginosa* heterologous Type II-A system utilized in (C), where phage is titrated in ten-fold dilutions (black circles) on a lawn of *P. aeruginosa* (white background) expressing the indicated anti-CRISPR proteins and low (0.001%), intermediate (0.01%), or high (0.1% arabinose) levels of SpyCas9–sgRNA programmed to target phage DNA. (D) In vitro DNA cleavage inhibition assay shows AcrIIA2b.3 has the least temperature-dependent inhibitory activity.

Cas9–sgRNA function was subsequently assayed at three different levels of induction, revealing an increase in phage targeting as the effector concentration increased (Figure 4C). The expression of AcrIIA4 yields robust inhibition of Cas9-based phage targeting at all induction levels of Cas9, while AcrIIA2 provides limited activity under the strongest Cas9 induction conditions but functioned well at lower levels of Cas9 (Figure 4C). Given the observed inefficiency of AcrIIA2 compared to AcrIIA4 in previous *E. coli* and human cell experiments (Rauch et al., 2017), it seems that this high level of Cas9–sgRNA induction mimics those experiments. Interestingly, two homologs from the AcrIIA2b family (AcrIIA2b.1 and AcrIIA2b.3) showed significantly stronger inhibition of SpyCas9 activity in comparison to AcrIIA2a, with AcrIIA2b.3 performing as well as AcrIIA4 (Figure 4C). Of note, expression of a protein from the AcrIIA2c family was toxic in *P. aeruginosa* and thus not pursued further. AcrIIA2b.1 did not express in *E. coli* and therefore we chose AcrIIA2b.3 (henceforth referred to as AcrIIA2b) for further biochemical analysis.

AcrIIA2b protein inactivates SpyCas9 function in a similar fashion to AcrIIA2 through direct interaction with sgRNA-loaded SpyCas9 (Figure S1D-1F). We next examined the extent to which AcrIIA2b can suppress SpyCas9-mediated DNA cleavage *in vitro*. In excellent agreement with our *in vivo* results, AcrIIA2b displays more robust inhibitory capacity against SpyCas9 activity than AcrIIA2 at both room temperature and body temperature (Figure 4D). Importantly, the inhibition exhibited by AcrIIA2b at all tested conditions was similar, if not better, than AcrIIA4 (Figure 4D). Competitive binding assays showed that AcrIIA2b competes with target DNA for binding to SpyCas9 when added to the enzyme simultaneously (Figure S10A), while preincubation of SpyCas9 with target DNA prevents engagement with AcrIIA2b (Figure S10C). Notably, the AcrIIA2b protein shows a greater affinity for SpyCas9 (K_D,app_=230 nM) compared to AcrIIA2 (K_D,app_=512 nM), whereas AcrIIA4 shows a comparable affinity (K_D,app_=78 nM) to target DNA (K_D,app_=26 nM) (Figure S10B). Taken together, these biochemical data indicate that AcrIIA2b recognizes the same binding site as AcrIIA2 in the SpyCas9–sgRNA binary complex and thereby prevents target DNA recognition and cleavage.

### Structure of AcrIIA2b-bound SpyCas9 reveals a distinct interaction network for PAM-recognition interference

To determine the underlying structural basis for AcrIIA2b-mediated SpyCas9 inhibition, we determined the cryo-EM structure of the SpyCas9–sgRNA complex bound to AcrII2b at ∼3.9 Å resolution (Figures 5A–B and S4B). In the SpyCas9–sgRNA–AcrIIA2b complex, AcrIIA2b is positioned in the PAM duplex DNA binding cleft, between the NUC and REC lobes. It occupies the same position as that occupied by AcrIIA2 (Figures 5C and S9B), although AcrIIA2b buries a larger solvent-accessible surface area (∼2,407Å^2^) upon ternary complex formation relative to AcrIIA2–Cas9 interface (∼1,803Å^2^) (Figure S7A, D). Despite a large sequence disparity, AcrIIA2b shares the same fold (α1α2β1β2β3α3β4) as AcrIIA2, and its overall structure is similar (Figures S7C, S7F, and S9C). Divergent loop regions form the binding crevice (Figure S7B, S7E, and S9C). Additionally, the SpyCas9 conformation observed in the AcrIIA2b-bound structure is nearly identical to that observed in the AcrIIA2-bound structure (Figure 5C), except that the Hel-II domain within the REC lobe undergoes an intermediate conformational change (Figure S6E–F).

**Figure 5.**
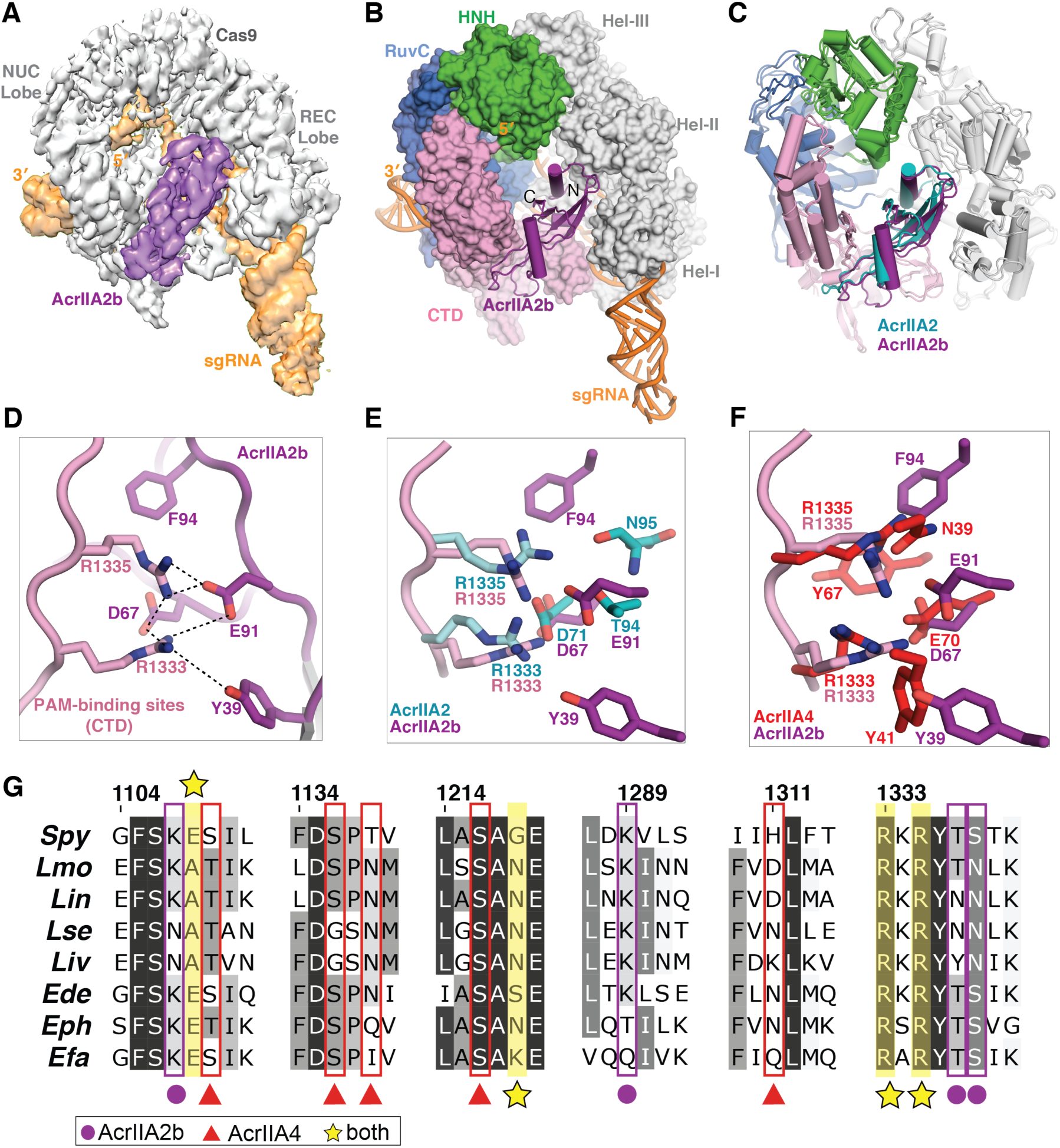
AcrIIA2b bears a strikingly similar interaction network to AcrIIA4 for PAM-recognition interference. (A) Cryo-EM reconstruction of the AcrIIA2b-bound SpyCas9. (B) The atomic model of SpyCas9–sgRNA–ArcrIIA2b. AcrIIA2b (purple) and sgRNA (orange) are shown in ribbon diagram and SpyCas9 is displayed as surface representation. (C) Structural comparison of AcrIIA2- and AcrIIA2b-bound SpyCas9–sgRNA complex structures. (D) Zoomed-in view of AcrIIA2b–SpyCas9 interaction at the PAM-recognition site. (E-F) Overlay of AcrIIA2b PAM-interference pocket with that of AcrIIA2 (E) and that of AcrIIA4 (F). (G) Alignment of Cas9 protein sequences denoting key residues that interact with the indicated anti-CRISPR proteins. Abbreviations: *Spy, Streptococcus pyogenes; Lmo, Listeria monocytogenes; Lin, Listeria innocua; Lse, Listeria seeligeri; Liv, Listeria ivanovii; Ede, Enterococcus devriesei; Eph, Enterococcus phoeniculicola; Efa, Enterococcus faecalis*.

Although AcrIIA2b utilizes a nearly identical binding mode to that deployed by AcrIIA2 to inhibit SpyCas9 enzymatic activity, a closer inspection of the AcrIIA2b binding crevice near the PAM-interacting site revealed a distinct local environment for interference with PAM recognition. Specifically, the two hydrophobic aromatic amino acid residues (Y39 and F94 in AcrIIA2b) that are engaged in hydrophilic and hydrophobic interactions with SpyCas9’s PAM-interacting residues are either not present or not involved in interfering with PAM recognition by AcrIIA2 (Figure 5E). Interestingly, this interaction pattern of AcrIIA2b bears a striking resemblance to that of AcrIIA4 (Figure 5F), although the two anti-CRISPRs are evolutionarily unrelated. Additionally, the residues that AcrIIA2b and AcrIIA4 bind are well evolutionarily conserved in Cas9 orthologues from across *Listeria* to closely related Cas9 orthologues from *Enterococcus* species (Figure 5G, S15 and S16). Based on these structural observations, we deduced that the presence of aromatic hydrophobic residues in the PAM-interacting site may contribute to the prominent *in vitro* and *in vivo* inhibitory function exerted by both AcrIIA2b and AcrIIA4, and that loss of this extensive hydrophobic interaction pattern in the AcrIIA2–Cas9 interface makes it less stable and therefore less effective for inhibiting SpyCas9 function.

To test this idea, we performed temperature- and urea-induced denaturation experiments on the SpyCas9–sgRNA–AcrIIA2b ternary complex and found that AcrIIA2b– SpyCas9 binding is more resistant to denaturation than is AcrIIA2a-SpyCas9 binding. To further assess the importance of individual residues for the inhibitory function of AcrIIA2b, we designed single and double amino acid substitutions and tested them using the *in vivo* anti-CRISPR activity assay in *P. aeruginosa* (Figures 4B). Substitution of Y39 or F94 for a small hydrophobic (Ala) residue reduced anti-CRISPR activity, whereas a Y39A/F94A double substitution decreased activity significantly, as observed with the D67A mutation of AcrIIA2b (Figures 6A and S17). *In vitro*, the Y39A/F94A double mutant showed a modest reduction in AcrIIA2b inhibition activity and more importantly, it displayed a more pronounced temperature-dependent anti-CRISPR activity, with the inhibition entirely ablated at body temperature (Figures 6B, S13B and S14). We concluded that the pattern of hydrophobic and hydrophilic residues surrounding the PAM-binding site is important to the function of anti-CRISPRs, and that anti-CRISPR stability and efficiency can be altered by large hydrophobic aromatic amino acid residues.

**Figure 6.**
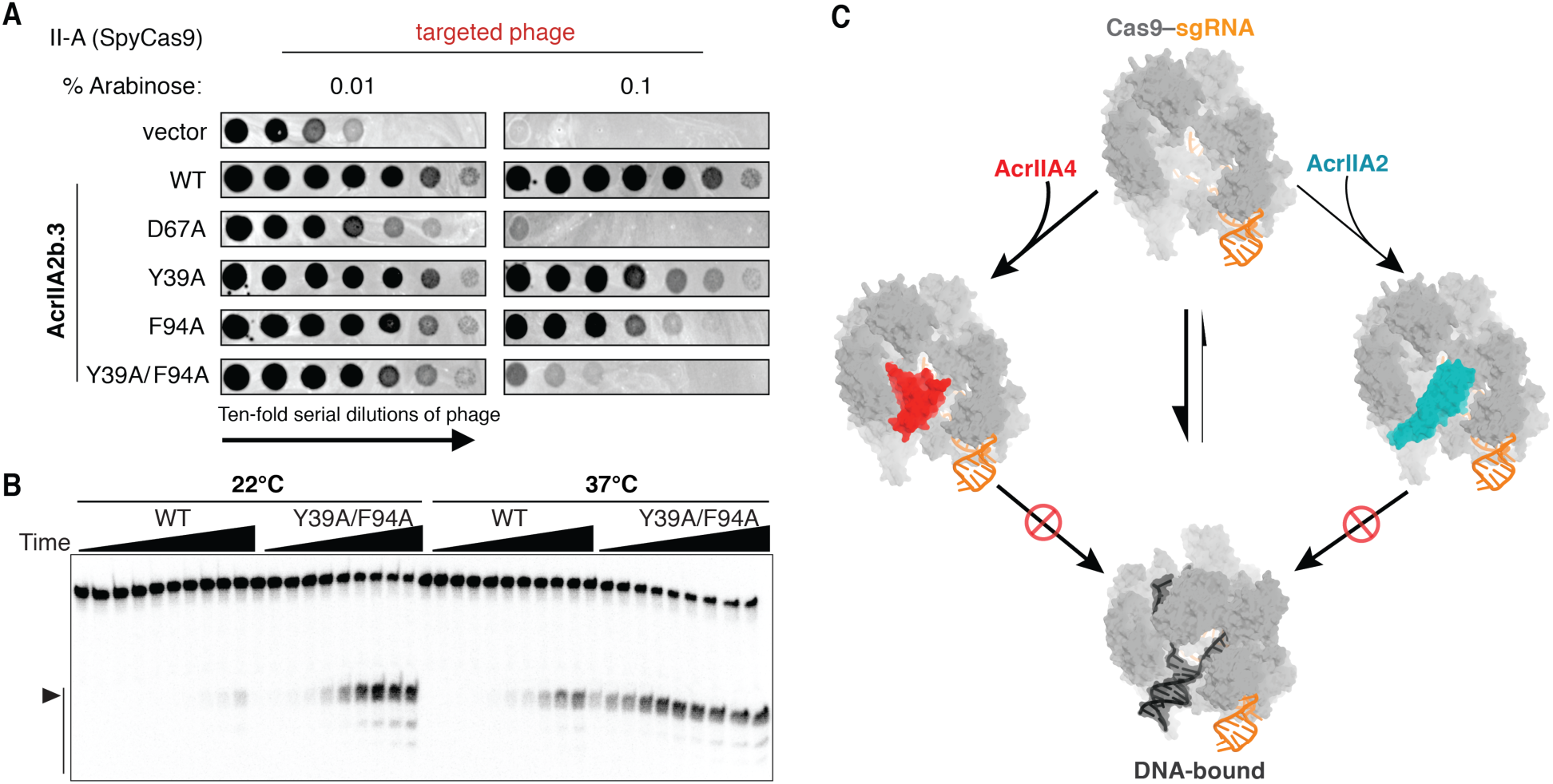
A convergent Cas9 inhibition mechanism shared by AcrIIA2 and AcrIIA4. (A) Plaquing of *P. aeruginosa* phage targeted by SpyCas9 in the presence of wildtype AcrIIA2b or the relevant point mutants. Double mutation of Y39A/F94A results in a large reduction of AcrIIA2b inhibition. (B) In vitro cleavage assay showing the double mutation is more vulnerable and sensitive to temperature. (C) Model of AcrIIA2 and AcrIIA4 convergent inhibition of SpyCas9’s target DNA binding and cleavage activities.

## DISCUSSION

### A convergent Cas9 inhibition mechanism between AcrIIA2 and AcrIIA4

Bacteriophage have evolved anti-CRISPRs to counteract bacterial CRISPR–Cas adaptive immunity. Anti-CRISPRs have diverse sequences ranging from 50 to 150 amino acids that lack similarity to previously reported structures (Bondy-Denomy, 2018; Pawluk et al., 2018). AcrIIA2 and AcrIIA4 represent the first example of anti-CRISPR proteins that can circumvent CRISPR-Cas targeting mediated by both type II-A *L. monocytogenes and S. pyogenes* Cas9 enzymes. Here we found that AcrIIA2 impedes PAM recognition and initial target DNA binding by occupying the PAM-interacting cleft on the sgRNA-loaded SpyCas9. In contrast to target DNA binding, which induces a large conformational change in the HNH and helical domains of SpyCas9, AcrIIA2 binding induces only a slight conformational change within SpyCas9.

Although AcrIIA2 and AcrIIA4 are unrelated anti-CRISPRs, their inhibition mechanisms both involve competitive inhibition of SpyCas9’s PAM recognition (Figure 6C). This convergent inhibition mechanism likely explains why AcrIIA2 and AcrIIA4 are mutually exclusive in prophage genomes (Rauch et al., 2017). It is also worth noting that, although many characterized anti-CRISPRs prevent target DNA binding (Bondy-Denomy et al., 2015), AcrIIA2 and AcrIIA4 are the only two examples to date that function by specifically blocking PAM recognition.

### Temperature-dependent anti-CRISPR activity and implications for genome editing control

*In vitro* assays revealed temperature-dependent anti-CRISPR inhibitory activity, with AcrIIA2 exhibiting a more pronounced temperature dependence compared to AcrIIA4. This observation may explain why AcrIIA2 can only partially inhibit SpyCas9 activity *in vivo*. The reduced temperature sensitivity of the AcrIIA2b homolog, which employs large aromatic residues to stabilize a favorable hydrophobic interaction on SpyCas9, also favors more robust Cas9 inhibition *in vivo*.

Previous studies have revealed that simultaneous delivery of AcrIIA4 with SpyCas9– sgRNA (RNP) complex inhibits Cas9-mediated gene targeting, and that proper timing of AcrIIA4 delivery can reduce off-target editing while retaining on-target editing levels (Shin et al., 2017). We expect that AcrIIA2 can be employed in a similar manner to AcrIIA4 for regulating Cas9 *in vivo* genome editing. The reduced temperature sensitivity of AcrIIA2b relative to AcrIIA2a supports the use of AcrIIA2b for future applications due to more robust inhibitory activity. Our studies provide a basis for structure-based design of efficacious peptide modulators or even small molecular inhibitors that specifically interfering with PAM recognition. With more structures of anti-CRISPRs solved in the near future, we hope to better understand their diverse inhibitory mechanisms, enriching our current knowledge of how anti-CRISPRs are used to leverage bacteria-phage arms race.

## AUTHOR CONTRIBUTIONS

J., J.B.D., and J.A.D designed experiments; F.J. and M. X. performed biochemistry experiments; F.J. and J.J. performed structural studies; B.A.O., J.D.B., B.J.R., and J.B.D. prepared strains and executed phage-plaquing experiments; J.A.D., J.B.D., and E.N. supervised experiments; F.J., J.B.D., and J.A.D wrote the manuscript with input from all authors.

## ACKNOWLEDGEMENTS

The Cryo-EM data was collected in the EM facility of University of California, Berkeley. We are thankful to D. Toso and P. Grob for expert electron microscopy assistance, and A. Chintangal and P. Tobias for computational support. We also thank members of the Bondy-Denomy, Doudna and Nogales labs for helpful discussions. F.J. is a Merck Fellow of the Damon Runyon Cancer Research (DRG-2201-14). J.B.D. is supported by the University of California San Francisco Program for Breakthrough in Biomedical Research, funded in part by the Sandler Foundation, and an NIH Office of the Director Early Independence Award (DP5-OD021344). J.A.D. and E.N. are Investigators of the Howard Hughes Medical Institute.

